## Supplementary Materials for "Combined responses of primary coral polyps and their algal endosymbionts to decreasing seawater pH"

Supplementary Methods

Figure S1. Experiment set up.

Figure S2. Larval survival and settlement.

Figure S3. Physiological parameters of coral host and algal symbiont.

Figure S4. Chlorophyll auto-fluorescence of live *S. pistillata* primary polyps and photosynthetic parameters.

Figure S5. Calcein Blue incorporation patterns and skeletal features measurements of *S. pistillata* primary polyps.

Figure S6. Nomenclature of structures in the primary polyps' skeleton and measurements of skeletal features.

Figure S7. Septa thickness measurements based on CT-scans.

Figure S8. Distribution of coral and symbiont differentially expressed genes.

Figure S9. Schematic summary of the response of coral primary polyps to OA.

Table S1. Parameters of the carbonate chemistry across experimental pH conditions.

Table S2. Count of HQ-reads mapped to the selected proteomes databases of *symbiodinium*/SAR and cnidarian/metazoan.

Table S3. Counts statistics of coral host and symbiont reads mapped to exons used in the DESeq2 DE analysis.

Table S4. Known coral biomineralization-related proteins.

Table S5. Photosynthetic parameters of algal endosymbiont.

Supplementary References

### Supplementary Methods

#### Sample collection and experimental design

Coral larvae were collected using larval traps (160 µm plankton net with a plastic collection container), that were placed on randomly selected colonies from the shallow reef (depth of 3-5 m) for several nights during April 2019, following expected peak releases (1). Actively swimming larvae were transported to a controlled environment aquarium system at the Leon H. Charney School of Marine Science at the University of Haifa. Per each pH treatment a total of 324 larvae were put in custom-made polypropylene plastic chambers consisting of a central cylinder sealed at the ends by 160 µm plankton netting. Settlement chambers were used in this experiment to avoid losing the larvae through the water recycle system of the aquariums before settlement. Similar settlement chambers were employed in previous works with coral larvae (2,3). Prior to the insertion of the larvae, the chambers were washed to remove any potential chemical released by the plastic, and they were left to soak in seawater for one week.

The settlement chambers (see Supplementary Fig. 1 for a graphic overview of the experimental set up) were placed in a system of 9 flow-through aquariums with artificial seawater (Red Sea Salt, Red Sea Ltd) replicating the spring northern Red Sea conditions: salinity of 40 g liter$^{-1}$, temperature of 22° C, and irradiance of 250 µmol photons m$^{-2}$ s$^{-1}$ on a 12-hour/12-hour photoperiod provided by a Mitras LX 7206 LED aquarium light system (GHL, Germany). Prior to the insertion of the settlement chambers, carbonate chemistry of seawater was manipulated in 6 of the experimental aquariums by injecting $CO_2$ to reduce the ambient pH (pH 8.2) and obtain the target values of pH 7.8 (3 aquariums) and pH 7.6 (3 aquariums). All aquariums per each pH treatment were connected and received water and light with the same conditions. For the entire duration of the experiment, temperature and salinity were monitored continuously by electrodes linked to a monitoring system (GHL, Germany) that controlled $CO_2$ injection, and they were also measured three times per day using a portable conductivity-TDS meter (GOnDO Electronic Co. Ltd., Taiwan).

Measurements of pH (NBS scale) were performed three times per day using a pH glass electrode (Metrohm, Switzerland). Buffer solutions (Rocker Scientific, Taiwan) were employed to perform a 3-points calibration (pH 4, pH 7 and pH 10) of the pH electrode. Measurements of total alkalinity (TA) were made once per day (triplicates) via titration with 0.1 N HCl containing 40.7 g NaCl liter$^{-1}$, using an automatic lkalinity titrator (855 Robotic Titrosampler, Metrohm, Switzerland) controlled by Tiamo Software (version 2.0, Metrohm, Switzerland). Automated titrations of 50 ml samples were performed. Parameters of seawater carbonate system were calculated from pH, TA, temperature, and salinity using the CO2SYS package (4) with constants from(5) as refit by(6) (Table S1).

During the experiment, all corals were fed once per day with 2 ml of concentrated planktonic suspension (Microvore, Brightwell R aquatics, United States).

Corals were left in their respective seawater treatments for 9 days prior collection. We ran the experiment for long enough to 1) analyze coral recruits with enough deposited skeleton to be able to detect changes at both macro- and micro-scale levels (7,8), 2) analyze coral recruits still at the primary polyp stage, before they could start to asexually form new polyps. At the end of the experiment, primary polyps were collected by gently pushing them out of the chambers using the tip of a razor blade.

**Coral survival and settlement**

After 9 days of experimental treatment exposure the numbers of settled primary polyps, still swimming larvae (not settled) and dead larvae (larvae dissolved during the experiment) were recorded. The number of dead larvae was measured by subtracting the number of swimming larvae + settled polyps from the total number of larvae that were placed at the beginning of the experiment in each pH condition. Numbers of settled primary polyps, swimming larvae and dead larvae are shown as percentages of the total number of larvae initially placed in each pH condition (N = 324, data from chambers in the same experimental pH were pooled together).

**Dark respiration rates measurements**

Randomly selected primary polyps (N = 6 primary polyps per pH treatment) were transferred directly from treatment aquaria to a 24-well glass microplate (Loligo systems, Denmark). Each microplate well (1700 µL) contained one single polyp. A one-hour dark incubation was performed in the same conditions of temperature and seawater chemistry as in experimental tanks. An optical fluorescence oxygen system (PreSens, Germany) was used to quantify oxygen consumption rates inside the sealed wells. Measurements were carried out for one hour in the dark. Data were recorded with MicroResp™

software (Loligo systems, Denmark). Before measurements, the oxygen sensors were calibrated against air-saturated seawater (100% oxygen) and a saturated solution of sodium sulfite (zero oxygen), as recommended by the user manual (Loligo systems, Denmark).

**Host protein, symbiotic algae isolation and pigment study**

Pools of 12 primary polyps per pH treatment were collected at the end of the experiment and were placed in 1.5 ml tubes (N = 3 tubes per pH treatment), snap frozen in liquid nitrogen and kept at -80°C. Samples were homogenized using an electrical homogenizer (HOG-160-1/2, MRC-labs, Israel), after the addition of 500 µl of filtered seawater. The homogenate was centrifuged at ×5000 g for 5 min at 4°C to separate the debris and the symbiont cells from the coral host tissue. Aliquots of 100 µL of the supernatant were used to determine the protein concentration of the coral host using the QPRO-BCA kit standard (Cyanagen, Italy) following the manufacturer protocol. A PerkinElmer (2300 EnSpire R, United States) plate reader was used to determine the total protein concentration at a 562 nm emission wavelength. Following a further centrifugation at ×5000 g for 5 min and the resuspension of the pellet, aliquots of 50 µL of the homogenate were used to determine the density of the symbiotic algae by fluorescent microscopic counts (Nikon Eclipse Ti, Japan), using a hemocytometer (BOECO, Germany). We used 4 replicates (1 $mm^2$ each) of cell counts per sample. Each replicate was photographed both in brightfield and in fluorescent light using 440 nm emission to identify chlorophyll to ensure counting of symbiont cells only. Cell counting was performed using NIS-Elements Advanced Research (version 4.50.00, Nikon, Japan). The resulting cell counts were normalized to average host protein concentration and to average host surface area per treatment (measured as described below). Chlorophyll-a (chl-a) concentrations were measured in 2 ml of homogenate that was incubated overnight with 90% cold acetone at 4°C. After incubation the samples were centrifuged at ×5000 g for 5 min at 4°C, and the optical density was measured in a 10 mm glass cuvette at wavelengths of 630, 647, 664 and 691 nm, using a NanoDrop (Thermo-Fisher, United States). The light absorbance results were used to calculate the chl-a concentration based on a previously described method (9). The resulting concentrations were normalized to algal cells number and to average primary polyp surface area per pH treatment.

**Fluorescence microscopy measurements of primary polyps**

Live coral primary polyps (N = 3 polyps per pH condition) were imaged using an Inverted Phase Contrast Fluorescent Microscope (Nikon Eclipse Ti, United States), immediately after the experimental exposure. Each sample was observed with a DS-Ri2 camera using a red fluorescence channel (emission 590 nm). Exposure and gain settings were kept constant between the samples (400 ms exposure and 1.2 gain). All images were acquired with the Nikon Nis-Elements software (Nikon Instruments, United States). Mean chlorophyll auto-fluorescence intensities were determined, after removing the background signal, using the mean gray value in FIJI (10), delimiting and selecting the area within the primary polyps edges (11).

The fluorescent dye calcein, which binds divalent ions (such as $Ca^{2+}$), has been extensively used to study coral calcification (12). A portion of the live primary polyps (N = 3 polyps per pH condition) was placed in Petri dishes (60 mm) and incubated for 2 hours in 2.6 μM Calcein Blue (Sigma 54375-47-2) dissolved in their respective experimental seawater. The polyps were then rinsed for 30 minutes in seawater from their correspondent experimental tank before observation. Primary polyps were imaged using an Inverted Phase Contrast Fluorescent Microscope (Nikon Eclipse Ti, United States); each sample was observed with a DS-Ri2 camera using the blue fluorescence filter DAPI (emission 420 nm). In order to measure the blue autofluorescence signal of the skeleton, primary polyps that were not incubated with Calcein Blue were imaged before and after removing the tissue. The tissue of all polyps (incubated and non-incubated with Calcein) was removed by soaking the polyps in 1% sodium hypochlorite (NaClO), and then the skeleton was rinsed with distilled water. All images were acquired with the Nikon Nis-Elements software (Nikon Instruments, United States). Images of the skeleton were normalized so that (i) background = 0, (ii) septa intensity is 100 %, and (iii) autofluorescence of the skeleton (without calcein) = 0. Mean Calcein Blue fluorescence intensities were determined using the mean gray value in FIJI (10), delimiting and selecting the area within the primary polyps edges (11). Images of live primary polyps stained with calcein were normalized so that (i) background = 0, (ii) mean intensity of the calyx (cavity hosting the live coral) is 100 %, and (iii) autofluorescence of the skeleton + tissue (without calcein) = 0.

**Photosynthetic parameters assessment**

Randomly selected primary polyps (N = 9 per pH condition) were dark-incubated for 20 minutes in the same conditions of temperature and seawater chemistry as in experimental tanks. A MAXI imaging Pulse Amplitude Modulation (iPAM) fluorometer (Waltz GmbH, Germany) was used to measure photosystem II (PSII) maximal quantum yield ($F_v/F_m$), non-photochemical quenching (NPQ) and relative electron transport rate (rETR) of the algal endosymbiont. All polyps were measured with an actinic light saturation pulse (2700 μmol photons $m^{-2}$ $s^{-1}$) after incubation at increasing light intensities (0, 1, 11, 21, 56, 81, 111, 146, 231, 336 and 461 μmol photons $m^{-2}$ $s^{-1}$). In each pH condition, the initial slope (α), relative maximal electron transport rate ($rETR_{max}$), and minimal photoinhibition point ($E_k$) were calculated from a fitted double exponential decay function following (13).

**Quantification of primary polyps' size and skeletal thickness**

Randomly selected primary polyps (N = 6 polyps per pH condition) were imaged using an Inverted Phase Contrast Fluorescent Microscope (Nikon Eclipse Ti, United States) under a 4x magnification. The planar footprint area of the primary polyps was calculated using the measurement tool of the Nikon Nis-Elements software (Nikon Instruments, United States). These measurements (Fig. S4D, no significant difference among pH conditions) were employed to normalize the data of the physiological parameters (Fig. S3B-C, E).

A laboratory micro-CT (Skyscan1172, Bruker micro-CT, Belgium) was used to image the primary polyps using source power settings of 80 kV and 124 μA and an image pixel size of 5 μm. Scan settings included a 0.5 mm thick aluminum filter, 360 rotation scan averaging of 3 radiographs per step, 0.8 s exposure times. Data were reconstructed using the manufacturer package nRecon (v 1.7.4.2, Brucker micro-CT, Belgium) with moderate ring and beam hardening correction settings (10 and 50%, respectively). Data were viewed in 3D using CTvox (v 3.0, Brucker-microCT, Belgium) and analyzed as a stack in FIJI (10). Thresholding on the z-stack was performed using the Otsu approach (14). Areas of interest (skeletal septa, see Fig. S7 for reference) were selected along the z-stack, and the voxel counter plug-in for FIJI (10) was used to determine the volume (thickness) of those areas (N = 6 septa per each polyp).

**Skeletal micromorphological analysis**

A portion of the primary polyps (N = 6 polyps per pH condition) was randomly collected at the end of the experiment and was placed in small Petri dishes (35 mm). The tissue of the polyps was removed by immersing the dish in 1% sodium hypochlorite (NaClO) for 10 minutes as formerly described (15). Thereafter, samples were rinsed with DI water followed by drying overnight. The polyps were vacuum coated with gold (for conductivity) prior to examination under a ZEISS SigmaTM SEM (Germany), by using a SE2 detector (8-10 kV, WD = 9-10 mm) and an in-lens detector (2 kV, WD = 1.5-2.5 mm). The area of the calyx and the crown area of the primary polyps were measured with FIJI delimiting and selecting the areas of interest in the SEM images (shaded areas in Fig. 2). The number of Rapid Accretion Deposits (RADs)(globular elements on the spine tips) was calculated from 3 randomly selected primary polyps per pH condition (3 spines per polyp, only spines located on the septa were analyzed) using the SEM images. The ratio was taken between the number of clearly visible RADs to surface area of the base of the spine, calculated with FIJI (10). Both measurements were taken along the spine axis. Thickness of the septa in the SEM images were measured from 3 randomly selected primary polyps per pH condition (3 septa per polyp, 3 measurements along the longitudinal axis of each septum), using FIJI (10). A description of the areas analyzed in the SEM images can be found in Fig. S5.

**RNA extraction, processing and sequencing**

Per each pH condition, primary polyps were collected at the end of the experiment and were placed in 1.5 ml tubes (3 tubes per pH, pools of 20-35 polyps per tube) containing 500 µl TRI Reagent Solution (Sigma-Aldrich, United Sates), snap frozen in liquid nitrogen and kept at -80°C prior to RNA extraction. Total RNA was extracted using the Invitrogen PureLink RNA micro kit (Thermo Fisher Scientific, United States) according to the manufacturer's protocol. DNAase treatment was performed within the RNA extraction procedures according to the manufacturers' instructions (Thermo Fisher Scientific, United States). RNA concentration was confirmed using a NanoDrop 2000 (Thermo Fisher Scientific, United States) and quality was tested on a TapeStation (Agilent Technologies, United States). Three independent samples were generated for each pH treatment. Sequencing libraries were prepared using a standard mRNA-Seq protocol (GENEWIZ, Germany). The polyA fraction (mRNA) was purified from 500 ng of total RNA following by fragmentation and the generation of double-stranded cDNA.

Then end repair, A base addition, adapter ligation and PCR amplification steps were performed. 150 bp paired-end reads were sequenced on an Illumina HiSeq High Output.

**Host and symbiont RNA-Seq differential expression analysis**

Standard RNA-Seq quality filtering was conducted as described previously (16): raw-reads were adapter trimmed using Cutadapt (17) and quality filtered using Trimmomatic (18). In order to detect the species origin of symbiont sequences, HQ-reads were blasted to selected proteomes databases of *symbiodinium*/SAR and cnidarian/metazoa, using Diamond (19). Top hits almost exclusively belonged to robust corals (mainly *S. pistillata*), and Symbiodiniaceae "clade A" species (mainly *Symbiodinium microadriaticum*) (Table S2). Reads were further aligned to the merged database of the host genome assembly (NCBI GCA_002571385.1) and the symbiont genome assembly (NCBI GCA_001939145.1) using STAR (20–22). For different samples, of all STAR concordantly-mapped reads, 92-97% were mapped to host exons, and 3.5-7.7% were mapped to symbiont exons (Table S3). DE analysis was conducted using Bioconductor DEseq (23), separately for the host and the symbiont genes, considering a single factor (pH) with three levels. The DEseq2 analysis was conducted based on host genes with ≥5 reads in ≥2 samples. For the symbiont, only genes whose 25% quantile of read count was at least 15 reads in ≥2 samples (with 12422 'survived' symbiont genes) were selected, since an incorrect between/within normalization in DESeq2 was found when using more liberal cutoffs for the tested *S. microadriaticum* data. Coral host and symbiont read normalization was indicated by obtaining typical distribution of genes in the DESeq2 MA plot (Fig. S8A-B). Among the genes that were found to be expressed in the host (23500 genes) and in the algal endosymbiont (12423 genes), we selected the ones with an adjusted P value < 0.05 and identified them as differentially expressed (DEGs).

**Host and symbiont functional enrichment**

Genes biological terms were assigned based on Uniprot *S. pistillata* (https://www.uniprot.org/), KEGG *S. pistillata* (https://www.kegg.jp/), *S. pistillata* Trinotate annotations (24), and Uniprot *S. microadriaticum* databases. Enrichment analysis was conducted in Bioconductor GOSeq, as described previously (16). Functional enrichment for *S. microadriaticum* was searched using the score-based tool GSEA (25), which may be more sensitive than p-value-based searches (like GOSeq) in cases of expected small non-significant changes in relatively large groups of genes. In GSEA, $\log_2$ fold changes were used as scores. Host and symbiont enriched terms with an adjusted P value < 0.05 were identified as significantly enriched.

As many enriched terms are functionally-related, for both the host and symbiont the biological terms were hierarchically clustered based on pairwise distances between groups of genes calculated according to Equation1

$$D = |xa \cap xb| / \text{minimum}(|a|, |b|) \qquad (1)$$

where $a$, $b$ are two sets of gene ids, and $x_a$ and $x_b$ are DE genes ids from $a$ and $b$, respectively. Terms trees were constructed using Bioconductor-ggtree (26). Heatmap representation of enrichment scores were based on 1) for the Bioconductor GOSeq data: the percentage of DEGs multiplied by -1 or by 1 for down or upregulated gene groups, respectively, and 2) for GSEA data: Normalized Enrichment Score (NES)(25).

**_S. pistillata_ biomineralization-related genes**

The expression of known coral biomineralization-related proteins (Table S4) from new coral skeletal proteome data (27) and from the literature (8,28–42) was tested against the differential expression results (DESeq2, as described above) among the comparisons pH 7.8 vs pH 8.2, pH 7.6 vs pH 8.2 and pH 7.6 vs pH 7.8. A BLAST search (https://blast.ncbi.nlm.nih.gov/Blast.cgi) was performed to identify these known coral biomineralization-related genes in the genome of *S. pistillata* (NCBI GCF_002571385.1). Fisher's exact test (P < 0.05) was used to determine the proportion of significantly up- vs down-regulated biomineralization genes.

**Statistical analysis**

Data of the larval survival and settlement were analyzed using Fisher's exact test. All other data were tested for normality (Shapiro-Wilk test) and homogeneity of variance (Brown–Forsythe test). If the assumptions of normality and homogeneity of variance were not met, data were transformed (square root). One-Way ANOVA was used for all measurements, followed by Tukey test for post hoc comparisons, in which significant groups have a value of P ≤ 0.05. Non-parametric equivalents of tests were used in cases where assumptions were violated despite transformations. For such cases, a Kruskal-Wallis test was used, followed by Dunn's test for multiple comparisons. All results are presented as mean ± standard error throughout. The GraphPad Prism software version 8.0.2 (GraphPad Inc.) was used to perform all the statistical tests.

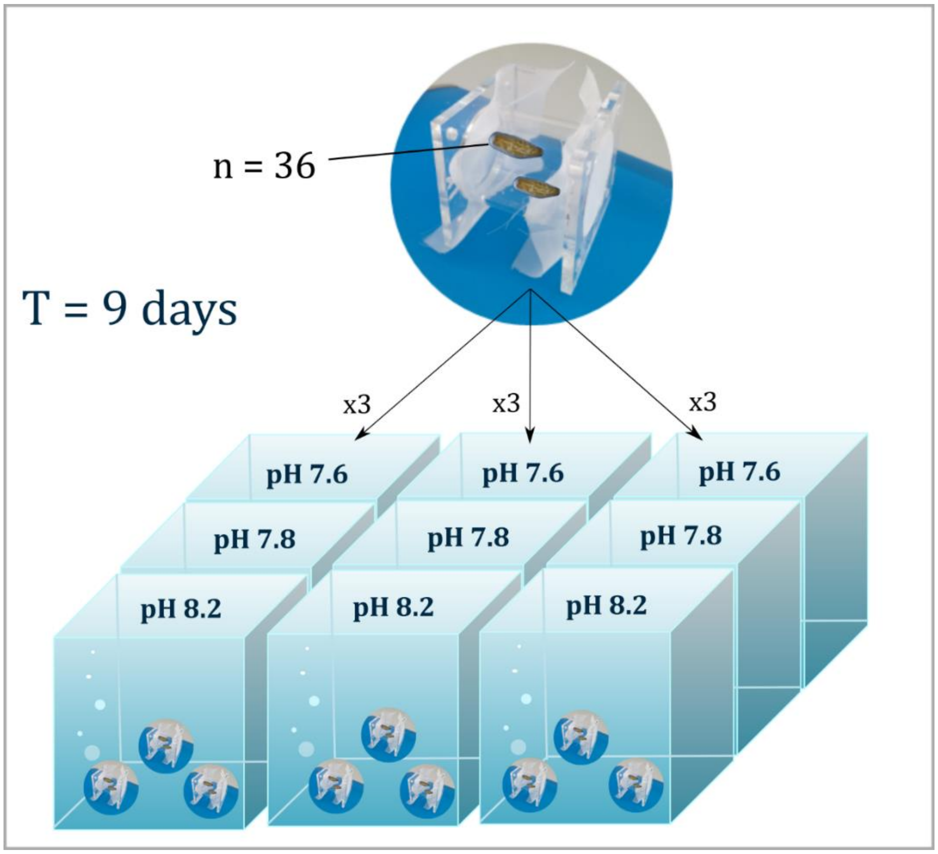


Figure S1. Experiment setup. Graphic summary of the experimental set up, from the allocation of coral larvae inside settlement chambers to the positioning of the chambers in the aquariums. T (Time), duration of the experiment.

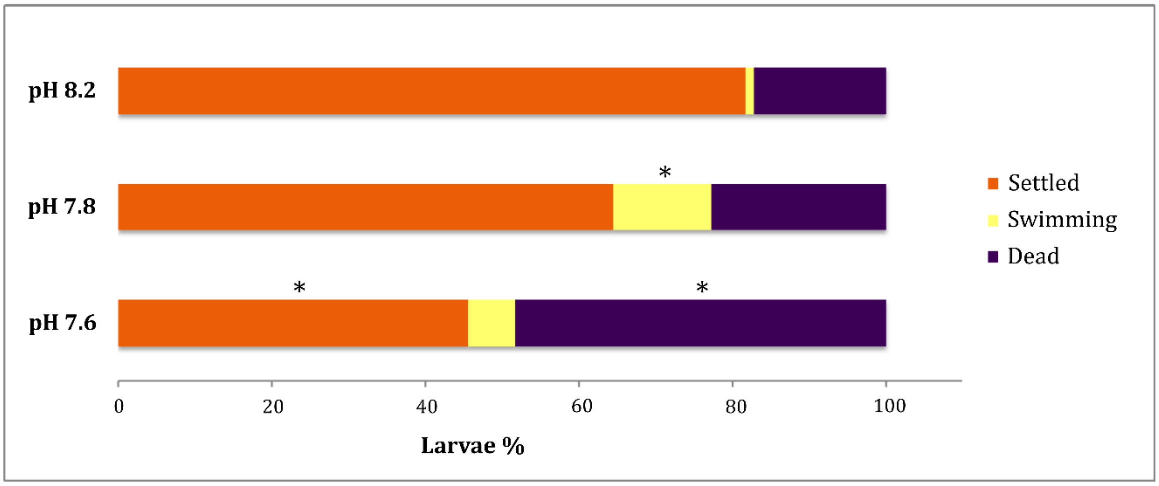


Figure S2. Larval survival and settlement. Stacked bar segments represent the mean percentages of larvae in each category (settled, swimming, dead) that were recorded at the end of the experiment in each pH condition, relative to the total number of larvae initially placed in each pH condition (N = 324 per pH). Orange indicates the percentage of larvae that metamorphosed and formed a primary polyp; yellow indicates the percentage of larvae that were still swimming and did not settle; purple indicates the percentage of larvae that died during the experiment. Asterisks (*) indicate statistical differences relative to the control pH 8.2 (Fisher's exact test, $P < 0.0001$).

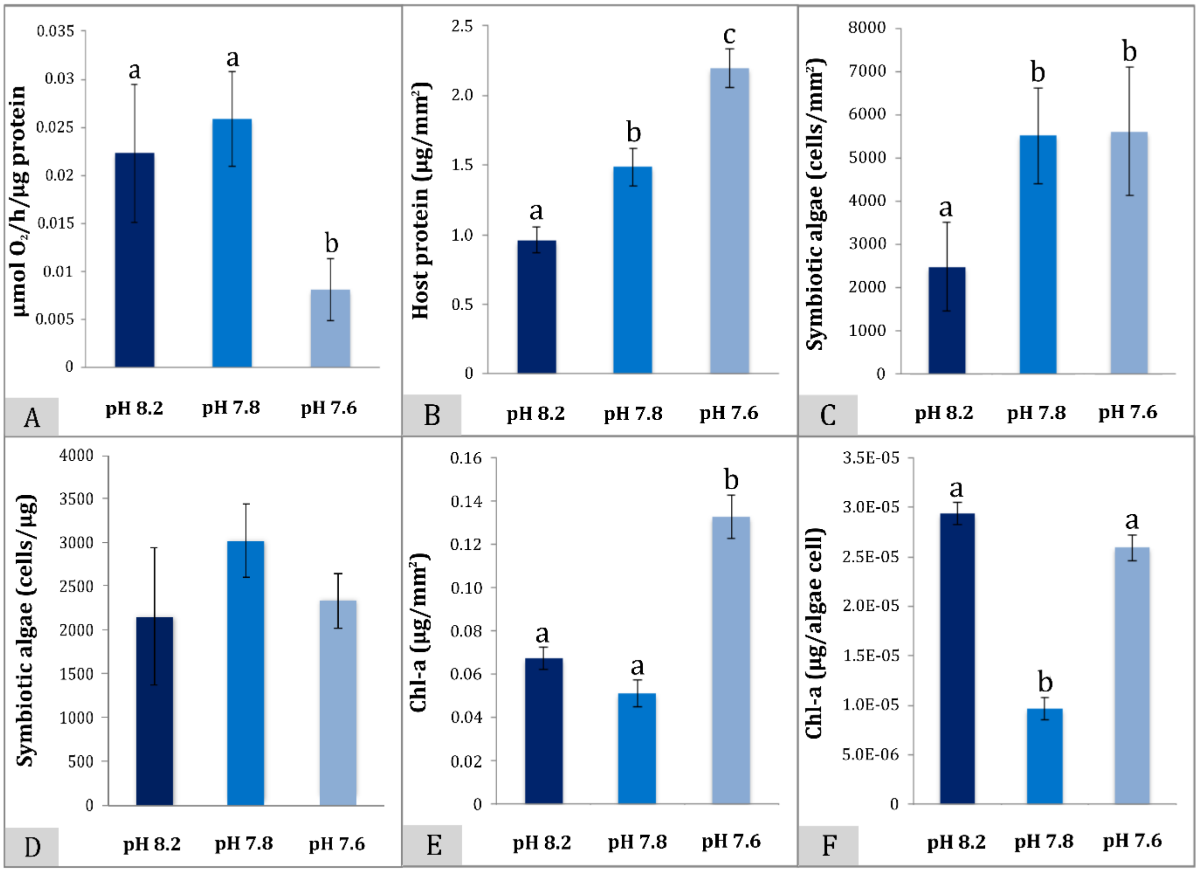


Figure S3. Physiological parameters of coral host and algal symbiont. (A) Primary polyps' respiration rates (means ± SEMs; N = 6 per pH condition), (B) coral host protein concentrations per surface area, (C) algae count per host surface area, (D) algae count per host protein, (E) chlorophyll-a concentrations per host surface area, and (F) chlorophyll-a concentrations per algae. Values are shown as means ± SEMs, N = 3 per pH condition. Different lowercase letters indicate statistical differences within the same panel ($P < 0.05$, One-Way ANOVA).

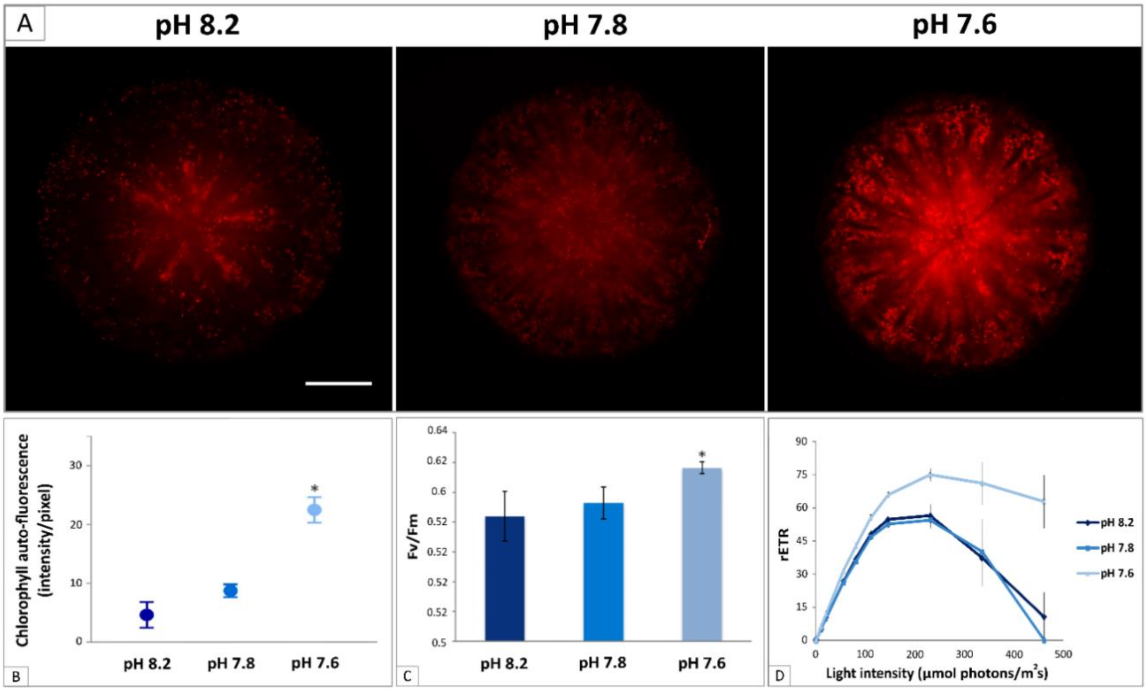


Figure S4. Chlorophyll auto-fluorescence of live *S. pistillata* primary polyps and photosynthetic parameters. (A) Microscopic images of live primary polyps. Red fluorescence indicates the chlorophyll auto-fluorescence of the symbiotic algae in the control pH 8.2, in the treatment pH 7.8, and in the treatment pH 7.6 (left to right). Microscope exposure and gain settings are identical in all three pH conditions. Magnification: 4x. Scale bar: 400 µm. (B) Chlorophyll auto-fluorescence intensity per pixel calculated from the microscopic images (means ± SEMs; N = 3 primary polyps per pH condition, only one polyp per pH is shown in panel A). (C) Algal maximum quantum yield ($F_v/F_m$); (D) algal relative electron transport rate (rETR). (B-D) Values are shown as means ± SEMs, N = 9 per pH condition. (B, C) Asterisks (*) within each graph indicate statistical differences relative to the control pH 8.2 (P < 0.05, One-Way ANOVA).

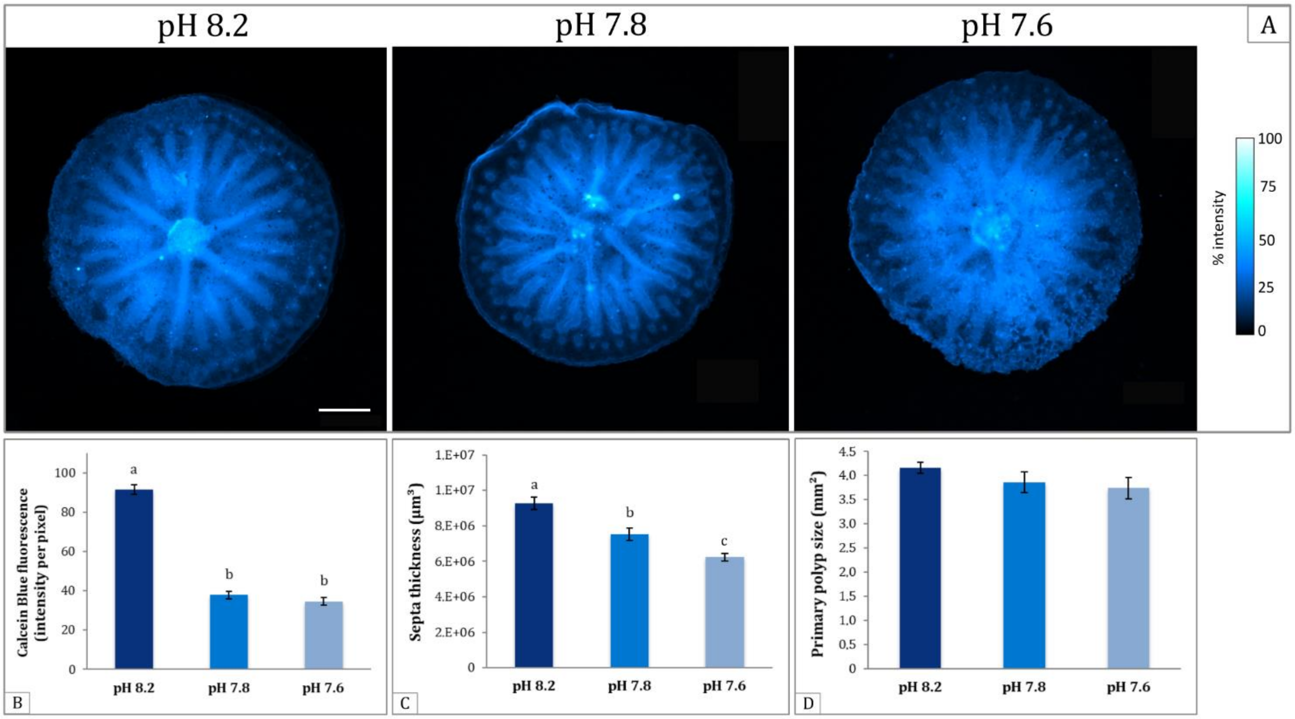


Figure S5. Calcein Blue incorporation patterns and skeletal features measurements of *S. pistillata* primary polyps. (A) Microscopic images of live primary polyps. Blue fluorescence indicates Calcein Blue fluorescence (normalized to maximum value and expressed as % intensity) in the live tissue and the skeleton in the polyps at the control pH 8.2, at the treatment pH 7.8, and at the treatment pH 7.6 (left to right). Magnification: 4x. Scale bar: 400 µm (B) Calcein Blue fluorescence intensity per pixel calculated from the microscopic fluorescence images of the primary polyps skeleton (without the tissue)(means ± SEMs; N = 3 primary polyps per pH condition, only one polyp per pH is shown in Figure 3). (C) Septa thickness (µm$^3$) measured from the CT-scans (means ± SEMs; N = 6). (D) Primary polyps mean planar area (mm$^2$). Absence of letters indicate no statistical difference among the three pH conditions (means ± SEMs; N = 6). (B, C) Different lowercase letters indicate statistical differences within each panel (P <0.0001, One-Way ANOVA).

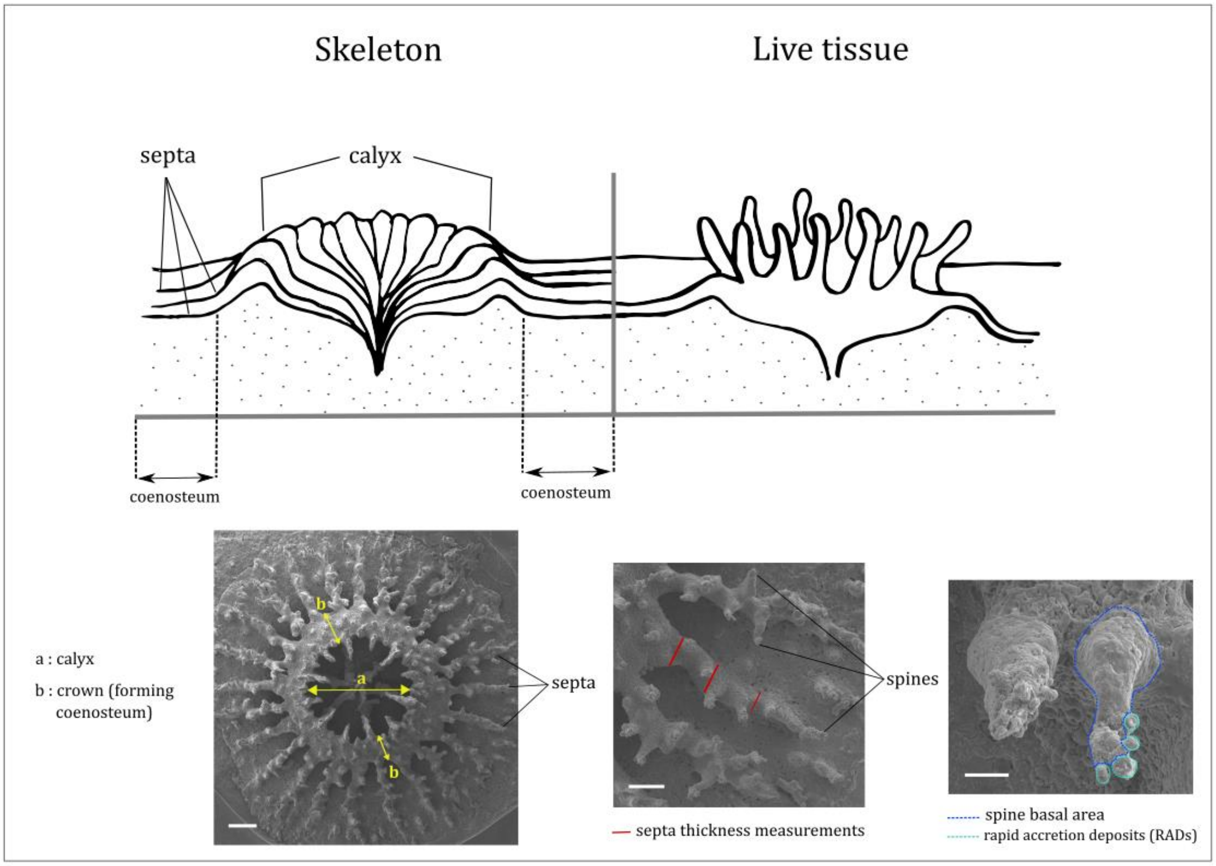


Figure S6. Nomenclature of structures in the primary polyps' skeleton and measurements of skeletal features. Upper section: Schematic representation of a coral live polyp (right) and of the underlying skeleton morphology (left). Skeletal features analyzed throughout this study are highlighted. Bottom section: Scanning electron micrographs showing (left to right) the skeleton of a primary polyp, enlargement of the septa and enlargements of spines on the septa. Yellow arrows indicating the calyx area ("a", cup-shaped opening hosting the coral) and the crown area ("b", forming coenosteum, here called "crown" for simplicity) that were measured in the primary polyps at the three levels of pH. Red segments indicating the three measurements of length taken along the spines to calculate septa thickness. Blue dotted line showing the basal area of the spine and cyan dotted line marking the RADs (rapid accretion deposits on the spines tip). The number of RADs on each spine was calculated per units of basal area of the spine. Scale bars (left to right): 200 µm, 100 µm,20 µm.

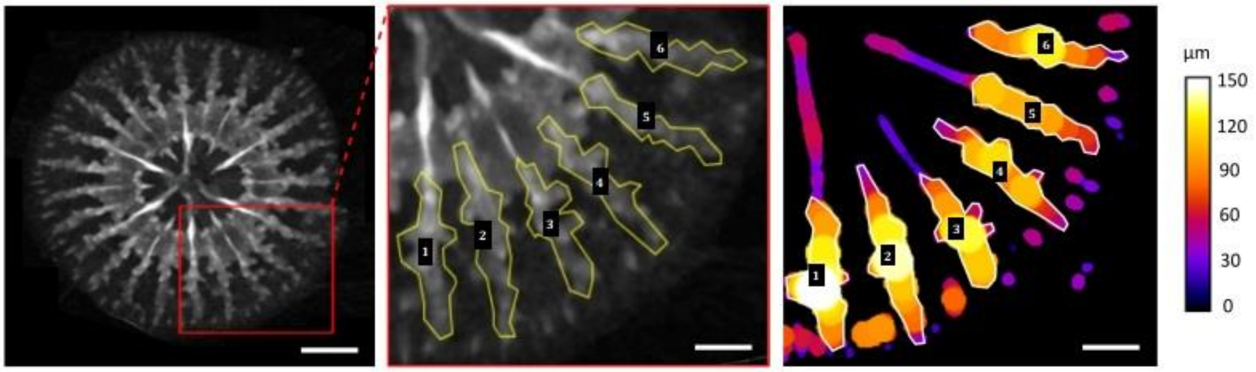


Figure S7. Septa thickness measurements based on CT-scans. Polygonal regions marking skeletal septa (N = 6) in the primary polyps that were measured to obtain and compare thickness values among pH conditions. Right picture showing the septa thickness distribution (µm) measured along the vertical axis. Scale bars: 400 µm (left picture), 200 µm (middle and right pictures).

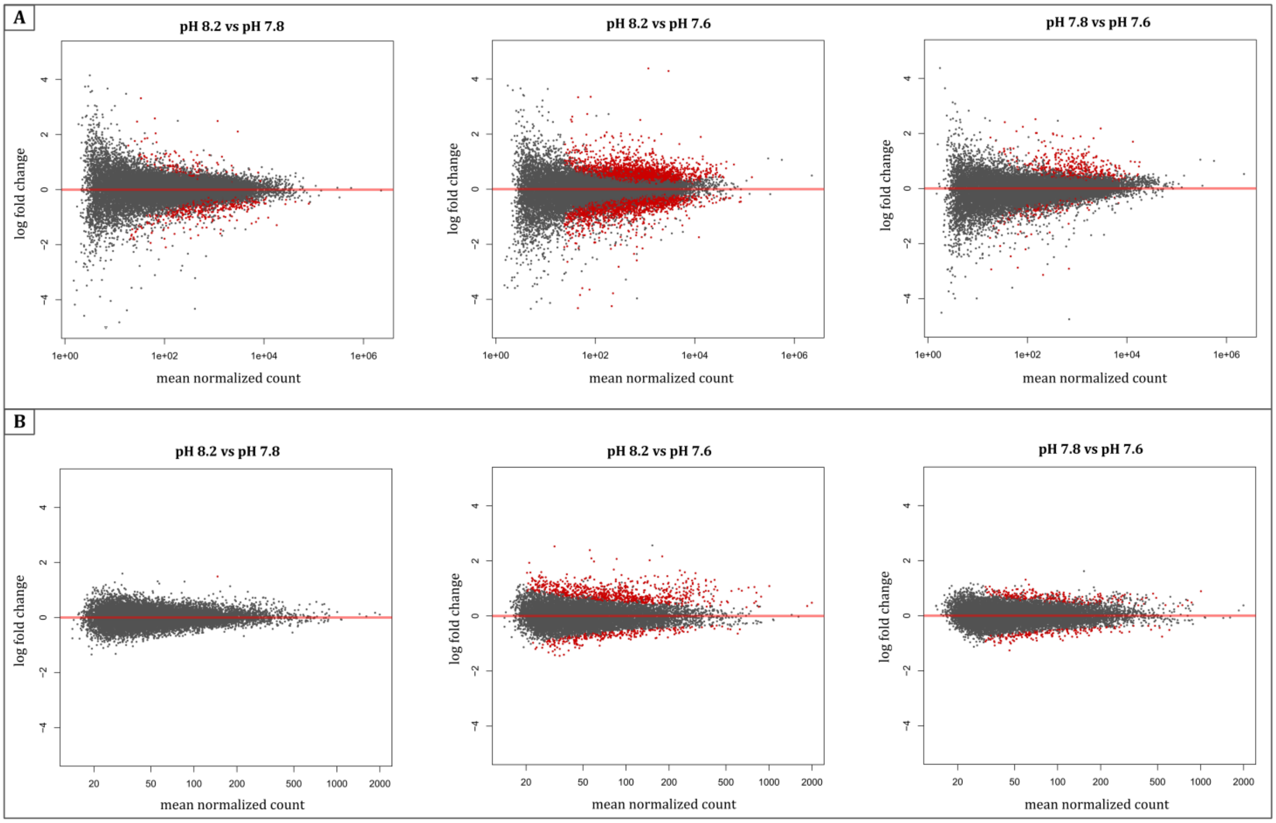


Figure S8. Distribution of coral and symbiont differentially expressed genes. (A) MA plots showing the coral reads normalization after the differential expression analysis performed using DESeq2. Red dots indicate coral expressed genes with a P value < 0.05. (B) MA plots showing the symbiont reads normalization after the differential expression analysis performed using DESeq2. Red dots indicate symbiont genes with a 25% quantile of read counts of at least 15.

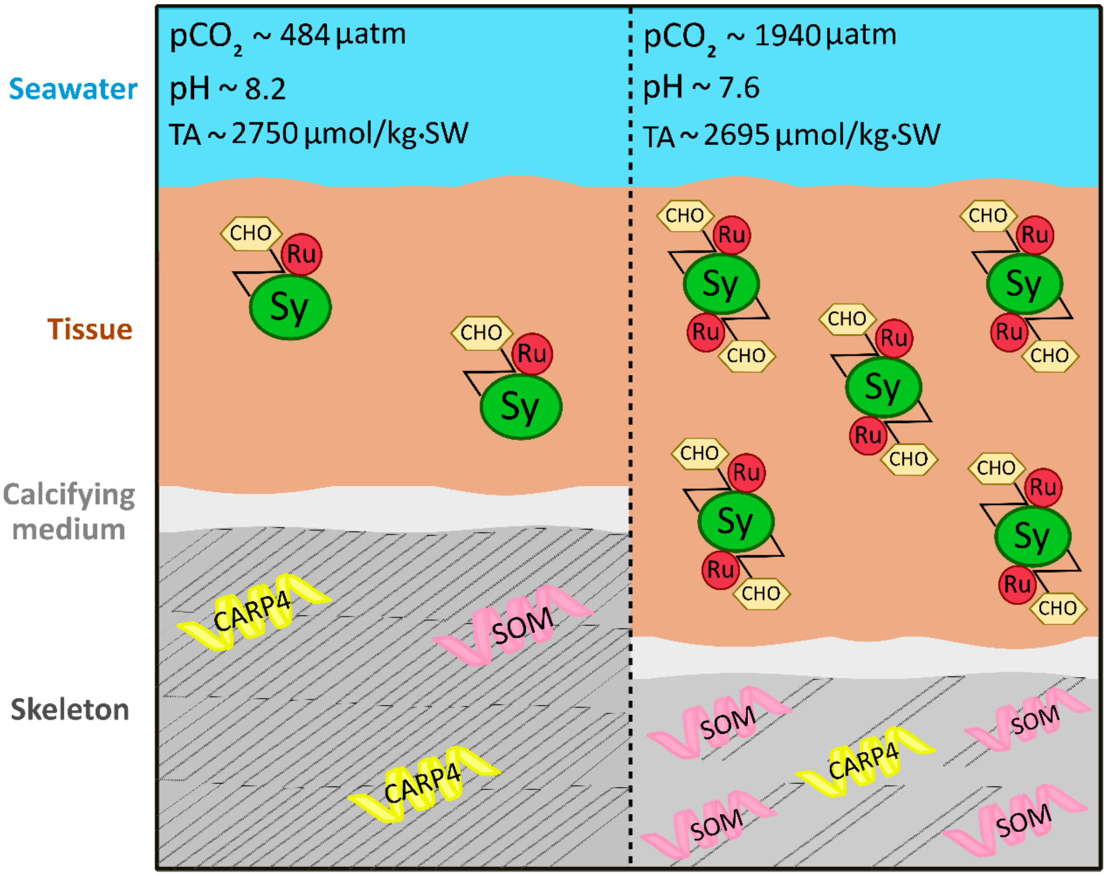


Figure S9. Schematic summary of the response of coral primary polyps to OA. Higher absorption of atmospheric $CO_2$ leads to a decrease of oceanic pH. Due to the increased energetic demands of the calcification process with OA, newly settled corals respond by building more porous and less developed skeletons, with a lower incorporation of biomineralization proteins that control the $CaCO_3$ deposition (CARP4). To counterbalance these changes and maintain overall growth, corals incorporate more skeletal organic matrix (SOM) proteins with structural functions and increase the amount of tissue covering the skeleton. With more available $CO_2$ stimulating the activity of the photosynthetic enzyme RuBisCO (Ru) and promoting symbiont photosynthesis and growth, the algae potentially enhanced the translocation of carbohydrates (CHO) to the host. Different textures of the skeleton indicate a more compact (left) versus a more porous skeleton (right). The “Z” symbol in correspondence of the algal cells (Sy) represents the photosynthetic process.

Table S1. Parameters of the carbonate chemistry across experimental pH conditions. Carbonate seawater chemistry parameters calculated from measured values of temperature (T), salinity (S), pH (NBS scale), and total alkalinity (TA). Measurements of pH, temperature and salinity were conducted three times a day, and measurement of total alkalinity were carried out once a day in triplicates for each experimental tank. Values are shown as means ± SD. DIC, dissolved inorganic carbon; Ωar, aragonite saturation state.

| Treatment | Tank | T (°C) | Salinity (‰) | pH | $CO_2$ (µmol/kg-SW) | $pCO_2$ (µatm) | $HCO_3^-$ (µmol/kg-SW) | $CO_3^{2-}$ (µmol/kg-SW) | DIC (µmol/kg-SW) | TA (µmol/kg-SW) | Ωar |
|---|---|---|---|---|---|---|---|---|---|---|---|
| **pH 8.2** | 1 | 22.11 | 39.7 | 8.16 | 14.3 | 482.3 | 2173.9 | 258.5 | 2446.7 | 2797.1 | 3.9 |
| | | ± 0.22 | ± 0.4 | ± 0.04 | ± 1.5 | ± 48.8 | ± 43 | ± 17.5 | ± 27.4 | ± 6.8 | ± 0.3 |
| | 2 | 22.13 | 39.7 | 8.16 | 14.2 | 476.2 | 2138.2 | 252.9 | 2405.3 | 2749.9 | 3.8 |
| | | ± 0.19 | ± 0.4 | ± 0.03 | ± 1.2 | 40.1 | ± 33.2 | ± 13.8 | ± 20.9 | ± 6.2 | ± 0.2 |
| | 3 | 22.28 | 39.8 | 8.14 | 14.6 | 494.0 | 2118.6 | 241.0 | 2374.2 | 2703.2 | 3.6 |
| | | ± 0.18 | ± 0.4 | ± 0.03 | ± 1.1 | 36.5 | ± 26.8 | ± 11.1 | ± 17.3 | ± 6.9 | ± 0.2 |
| **pH 7.8** | 4 | 22.44 | 39.7 | 7.79 | 37.1 | 12.56.3 | 2404.8 | 122.5 | 2564.4 | 2701.2 | 1.9 |
| | | ± 0.13 | ± 0.5 | ± 0.02 | ± 2.3 | 76.1 | ± 14.8 | ± 6.4 | ± 11.3 | ± 6.4 | ± 0.1 |
| | 5 | 22.39 | 39.7 | 7.80 | 36.6 | 1237.8 | 2396.9 | 124.1 | 2557.6 | 2697.3 | 1.9 |
| | | ± 0.13 | ± 0.5 | ± 0.04 | ± 3.9 | ± 130 | ± 27.9 | ± 11.1 | ± 21.3 | ± 8.9 | ± 0.2 |
| | 6 | 22.37 | 39.8 | 7.79 | 36.9 | 1249.4 | 2385.9 | 121.3 | 2544.10 | 2680.0 | 1.8 |
| | | ± 0.08 | ± 0.4 | ± 0.03 | ± 2.8 | ± 95.5 | ± 19.6 | ± 7.2 | ± 15.7 | ± 7.5 | ± 0.1 |
| **pH 7.6** | 7 | 22.07 | 39.6 | 7.62 | 58.7 | 1969.8 | 2509.2 | 84.1 | 2652.0 | 2712.5 | 1.3 |
| | | ± 0.29 | ± 0.5 | ± 0.04 | ± 5.9 | ± 198.1 | ± 18.7 | ± 7.2 | ± 17.5 | ± 6.7 | ± 0.1 |
| | 8 | 22.32 | 39.9 | 7.63 | 56.5 | 1910.3 | 2482.3 | 87.2 | 2625.9 | 2693.4 | 1.30 |
| | | ± 0.11 | ± 0.4 | ± 0.06 | ± 9.1 | ± 304.5 | ± 29.9 | ± 11.2 | ± 27.7 | ± 5.9 | ± 0.2 |
| | 9 | 22.27 | 39.7 | 7.62 | 57.4 | 1939.0 | 2476.00 | 84.1 | 2617.5 | 2679.7 | 1.3 |
| | | ± 0.09 | ± 0.4 | ± 0.04 | ± 5.7 | ± 190.8 | ± 19.8 | ± 7.2 | ± 18.5 | ± 6.5 | ± 0.1 |

Table S2. Count of HQ-reads mapped to the selected proteomes databases of *symbiodinium*/SAR and cnidarian/metazoan. Species origin of symbiont sequences detected in three replicates per pH condition by mapping the HQ-reads to selected proteomes databases of *symbiodinium*/SAR and cnidarian/metazoa, using Diamond. Blue and yellow rows indicate the top hits belonging to robust corals (mainly *S. pistillata*) and to Symbiodiniaceae "clade A" species (mainly *Symbiodinium microadriaticum*) respectively. Letters a, b and c indicate different replicates per pH condition.

| species | clade | pH7.6a | pH7.6b | pH7.6c | pH7.8a | pH7.8b | pH7.8c | pH8.2a | pH8.2b | pH8.2c |
|---|---|---|---|---|---|---|---|---|---|---|
| **CniStylophoraP** | other | 8468180 | 7667026 | 10073195 | 8180922 | 9540492 | 6796028 | 6588283 | 6175709 | 8860723 |
| **CniSeriatoporaSp** | other | 2335110 | 2372122 | 2734735 | 2386754 | 2598067 | 2927470 | 2007602 | 2548480 | 2389793 |
| **CniPocilloporaA** | other | 1504355 | 1347050 | 1727089 | 1463778 | 1701931 | 1310880 | 1175858 | 1127874 | 1554565 |
| **Symbiodinium Microadriaticum** | Symbiodinium | 871846 | 1238547 | 1742452 | 814225 | 1224972 | 828987 | 470002 | 282971 | 732855 |
| **Symbiodiniumn Clade A3** | Symbiodinium | 446606 | 638913 | 876443 | 422280 | 617842 | 437963 | 258236 | 167192 | 379470 |
| **clade A total** | | 1318452 | 1877460 | 2618895 | 1236505 | 1842814 | 1266950 | 728238 | 450163 | 1112325 |
| CniNematostellaV | other | 242199 | 478283 | 250750 | 361673 | 212470 | 1147198 | 396304 | 987401 | 195036 |
| CniFungiaSp | other | 333239 | 292275 | 358483 | 315748 | 352447 | 282039 | 264131 | 256155 | 340965 |
| CniGoniastreaA | other | 259435 | 231611 | 280604 | 251579 | 275550 | 233252 | 214431 | 211718 | 279162 |
| CniOrbicellaF | other | 211158 | 185080 | 230102 | 197790 | 224570 | 176166 | 169066 | 163180 | 215360 |
| CniPoritesL | other | 185356 | 158844 | 190282 | 176868 | 187711 | 159506 | 157696 | 153836 | 192764 |
| CniMontiporaC | other | 175927 | 157583 | 196391 | 170109 | 188073 | 149885 | 148600 | 137981 | 194056 |
| CniGalaxeaF | other | 171698 | 144083 | 172444 | 164212 | 176729 | 154938 | 141571 | 142072 | 173372 |
| SarFragilariopsisC | other | 63281 | 151030 | 62273 | 86670 | 41627 | 268269 | 144608 | 253601 | 51284 |
| CniAcroporaD | other | 122566 | 114217 | 139845 | 118879 | 135228 | 109875 | 100711 | 97161 | 128774 |
| CniAmplexidiscus | other | 104544 | 97013 | 112750 | 103118 | 111742 | 97189 | 87243 | 80225 | 110895 |
| DanioR | other | 57744 | 111308 | 58612 | 75976 | 51809 | 199131 | 95486 | 173314 | 60006 |
| CniDiscosoma | other | 92935 | 89624 | 101244 | 91563 | 99493 | 91108 | 79244 | 75363 | 96488 |
| SarEctocarpusS | other | 55822 | 99408 | 41006 | 67973 | 34491 | 189088 | 96098 | 169744 | 46638 |
| CniAcroporaM | other | 89528 | 82055 | 100357 | 87408 | 97648 | 75496 | 74667 | 68920 | 96466 |
| CniMorbakkaV | other | 44616 | 94346 | 47097 | 66510 | 38612 | 177911 | 82937 | 164307 | 42048 |
| CniAcroporaT | other | 83094 | 76837 | 95213 | 79905 | 91039 | 70894 | 68189 | 64013 | 87647 |
| CniHydraV | other | 32242 | 63410 | 32821 | 51414 | 26308 | 141262 | 51829 | 122293 | 29037 |
| MusM | other | 61358 | 65623 | 61970 | 64381 | 61389 | 66082 | 39789 | 39650 | 65847 |
| SarThalassiosiraO | other | 27229 | 61750 | 25418 | 34747 | 18247 | 81615 | 65174 | 83430 | 26361 |
| SymbiodiniumCladeB | Symbiodinium | 36422 | 57439 | 65190 | 38628 | 46506 | 39190 | 30378 | 17099 | 39577 |
| Human | other | 39629 | 40819 | 37563 | 38343 | 37123 | 36023 | 33077 | 27291 | 42827 |
| SarPhaeodactylumT | other | 31541 | 46063 | 29685 | 19620 | 9522 | 15626 | 68484 | 27434 | 29243 |
| StrongylocentrotusP | other | 31438 | 37468 | 31677 | 31755 | 31421 | 28899 | 33577 | 23699 | 40986 |
| AcanthasterP | other | 28074 | 35447 | 26236 | 29080 | 25477 | 28097 | 32195 | 25723 | 38938 |
| ChlVolvoxC | other | 17135 | 35197 | 14355 | 22867 | 12801 | 60581 | 28833 | 56145 | 16750 |
| SolanumL | other | 14285 | 37520 | 11305 | 20922 | 9365 | 59710 | 33900 | 55785 | 13889 |
| ChlamydomonasR | other | 17458 | 35047 | 15389 | 21912 | 14454 | 47846 | 28475 | 43486 | 17935 |
| SymbiodiniumSpCladeC | Symbiodinium | 23288 | 32838 | 37918 | 24140 | 28241 | 21980 | 18326 | 10455 | 25314 |
| DrosophilaM | other | 25450 | 29021 | 21819 | 24897 | 21466 | 22162 | 31646 | 19318 | 29161 |
| CniRenillaM | other | 23956 | 26236 | 24682 | 23464 | 22925 | 20278 | 25320 | 17887 | 28543 |
| alvTetrahymenaT | other | 10609 | 67186 | 6785 | 13969 | 5046 | 36880 | 35188 | 30153 | 10191 |
| SarNannochloropsisG | other | 11469 | 23327 | 10449 | 17337 | 8810 | 47580 | 21063 | 42164 | 8594 |
| CniDendronephthya | other | 19330 | 22914 | 17382 | 19533 | 16153 | 16793 | 23355 | 14986 | 24950 |
| SymbiodiniumGoreaui | Symbiodinium | 15873 | 26707 | 24578 | 17044 | 18292 | 15648 | 15188 | 7808 | 18030 |
| SarAureococcusA | other | 10823 | 25328 | 7017 | 15159 | 5827 | 32017 | 26087 | 26705 | 12110 |
| SymbiodiniumKawagutii | Symbiodinium | 12720 | 20530 | 20572 | 13711 | 15300 | 11914 | 11471 | 5528 | 14374 |
| PhyscomitrellaP | other | 10172 | 22160 | 6753 | 12610 | 5624 | 22275 | 20722 | 19855 | 10419 |
| alvBabesiaBi | other | 6307 | 16262 | 6447 | 9620 | 5174 | 28439 | 13369 | 24197 | 5431 |
| CniAureliaA | other | 11016 | 17367 | 10218 | 11725 | 9747 | 10831 | 13314 | 8600 | 14916 |
| SarThalassiosiraP | other | 8622 | 17780 | 6905 | 8236 | 2741 | 9206 | 24271 | 11138 | 9052 |
| AmborellaT | other | 5490 | 16473 | 2907 | 6401 | 2414 | 12732 | 18056 | 18797 | 6554 |
| BotrylloidesL | other | 9162 | 13134 | 6430 | 9094 | 6159 | 7736 | 12943 | 6010 | 15396 |
| MarchantaP | other | 6712 | 16623 | 3628 | 7170 | 2849 | 10531 | 17027 | 10766 | 6864 |
| alvParameciumT | other | 5557 | 36314 | 1724 | 5741 | 841 | 6429 | 19570 | 3427 | 5533 |
| CaenorhabditisE | other | 8088 | 14662 | 5244 | 8361 | 4682 | 6809 | 13003 | 4894 | 10930 |
| SelaginellaM | other | 5946 | 18681 | 2914 | 6690 | 2245 | 6266 | 15452 | 5465 | 6945 |
| ArabidopsisT | other | 7263 | 14644 | 3361 | 6992 | 3017 | 6570 | 13616 | 4704 | 7887 |
| KlebsormidiumN | other | 6081 | 12150 | 2506 | 6597 | 2338 | 5594 | 12943 | 4386 | 7420 |
| chlOstreococcusL | other | 5470 | 13074 | 4249 | 5189 | 3521 | 5885 | 9878 | 5075 | 5614 |
| ChlMonoraphidiumN | other | 5993 | 9408 | 2874 | 6242 | 3032 | 5496 | 10189 | 4379 | 7017 |
| CniThelohanellusK | other | 4083 | 7503 | 3312 | 4762 | 3470 | 9946 | 5883 | 7993 | 3671 |
| ChlAuxenochlorellaP | other | 4310 | 9590 | 2014 | 4861 | 1847 | 5444 | 11576 | 6028 | 5606 |
| chlOstreococcusT | other | 4972 | 7058 | 3585 | 4914 | 3390 | 4348 | 8035 | 3799 | 5718 |
| alvIchthyophthiriusM | other | 2671 | 21821 | 828 | 2845 | 450 | 4330 | 9797 | 1697 | 2557 |
| ChlMicromonasC | other | 4023 | 9445 | 1401 | 4136 | 1198 | 3585 | 9562 | 3303 | 5061 |
| alvPlasmodiumB | other | 3055 | 5131 | 1994 | 3435 | 2277 | 3396 | 3816 | 1611 | 2906 |
| alvPlasmodiumCh | other | 2898 | 4050 | 1133 | 3335 | 1277 | 3895 | 5393 | 1100 | 2956 |
| alvEimeriaN | other | 1808 | 6340 | 1161 | 2172 | 986 | 1656 | 4399 | 1171 | 2036 |
| alvBabesiaBo | other | 1489 | 4850 | 1063 | 1570 | 907 | 2129 | 3515 | 1977 | 1584 |
| alvEimeriaC | other | 1597 | 5579 | 645 | 1700 | 527 | 1357 | 3931 | 920 | 1985 |
| alvPlasmodiumCy | other | 1146 | 2713 | 673 | 1105 | 755 | 1012 | 2254 | 652 | 1089 |
| alvPlasmodiumCo | other | 956 | 2531 | 368 | 758 | 222 | 753 | 1940 | 478 | 870 |
| alvBabesiaO | other | 651 | 1965 | 294 | 609 | 259 | 606 | 1326 | 413 | 642 |
| alvBabesiaSp | other | 674 | 1744 | 238 | 603 | 175 | 510 | 1419 | 331 | 623 |
| alvEimeriaM | other | 653 | 1400 | 289 | 653 | 312 | 516 | 1255 | 363 | 667 |

Table S3. Counts statistics of coral host and symbiont reads mapped to exons used in the DESeq2 DE analysis. Count of STAR concordantly-mapped reads to the host genome assembly (NCBI GCA_002571385.1) and to the symbiont genome assembly (NCBI GCA_001939145.1), and relative percentages of the total reads mapped to the host exons and to the symbiont exons. Total count indicates the total number of concordantly-mapped genes for both *S. pistillata* and *S. microadriaticum*. Letters a, b and c indicate different replicates per pH condition.

| species | pH7.6_a | pH7.6_b | pH7.6_c | pH7.8_a | pH7.8_b | pH7.8_c | pH8.2_a | pH8.2_b | pH8.2_c |
|---|---|---|---|---|---|---|---|---|---|
| *S. pistillata* count | 25345107 | 22485863 | 27379682 | 25568291 | 27761001 | 24759788 | 20318634 | 22256167 | 26057103 |
| *S. microadriaticum* count | 924626 | 1292424 | 1824936 | 862279 | 1294561 | 886265 | 496498 | 300094 | 771524 |
| *S. pistillata* % | 96.5% | 94.6% | 93.8% | 96.7% | 95.5% | 96.5% | 97.6% | 98.7% | 97.1% |
| *S. microadriaticum* % | 3.5% | 5.4% | 6.2% | 3.3% | 4.5% | 3.5% | 2.4% | 1.3% | 2.9% |
| total | 26269733 | 23778287 | 29204618 | 26430570 | 29055562 | 25646053 | 20815132 | 22556261 | 26828627 |

Table S4. Known coral biomineralization-related proteins. Accession numbers/gene IDs of know biomineralization-related proteins from new coral skeletal proteome data (27) and from the literature (8,28–42). The first reference in the References column is the one related to the accession number/gene ID, all other references relate to studies where that specific gene/protein was previously detected/immunolocalized (chronological order).

| Accession number/Gene ID | Definition | References |
|---|---|---|
| MG182344.1; MG182345.1 | Acropora yongei Na+/Ca2+ exchanger | Barron et al., 2018 |
| EU532164.1 | carbonic anhydrase 2 | Bertucci et al., 2011 |
| AGG36361.1 | Annotated: Protoacadherin (PC4) | Drake et al., 2013 |
| P12 g2385 | Stylophora pistillata hypothetical protein | Drake et al., 2013 |
| P13 g6918 | Sushi domain-containing | Drake et al., 2013 |
| P14 g9951 | clone g9951 alpha collagen-like protein gene | Drake et al., 2013 |
| P16 g11702 | Stylophora pistillata clone g11702 hypothetical protein gene | Drake et al., 2013 |
| P18 g810 | Stylophora pistillata clone g810 alpha collagen-like protein gene | Drake et al., 2013 |
| P19g20041 | Contactin-associated protein | Drake et al., 2013 |
| P20 g6066 | MAM domain anchor protein | Drake et al., 2013 |
| P21 g18277 | Zona pellucida | Drake et al., 2013 |
| P22 g19762 | Stylophora pistillata clone g19762 hypothetical protein gene | Drake et al., 2013 |
| P23 g1057 | Protocadherin | Drake et al., 2013 |
| P24 g15888 | clone g15888 vitellogenin-like protein gene | Drake et al., 2013 |
| P26 g1441 | clone g1441 vitellogenin-like protein gene | Drake et al., 2013 |
| P27 g18472 | Integrin - alpha | Drake et al., 2013 |
| P28 g11651 | Late embryogenesis protein | Drake et al., 2013 |
| P3 g12510 | Thrombospondin | Drake et al., 2013 |
| P31 g20420 | Neurexin | Drake et al., 2013 |
| P32 g5540 | Kielin/chordin llke | Drake et al., 2013 |
| P33 g8985 | Flagellar associated protein | Drake et al., 2013 |
| P34 g1714 | MAM/LDL receptor domain containing protein | Drake et al., 2013 |
| P36 g13890 | Zonadhesion-like precursor | Drake et al., 2013 |
| P4 g9861 | Viral inclusion protein | Drake et al., 2013 |
| P5 g11674 | Hemicentin | Drake et al., 2013 |
| P8 g9654 | Major yolk protein | Drake et al., 2013 |
| P9 g10811 ; P1 g11108 ; P10 g11107 | Protocadherin fat-like | Drake et al., 2013 |
| JN631095.1 | Acropora millepora clone B26 hypothetical protein p251_4 | Hayward et al., 2011 |
| Gene g29033.t1 | Annotated: Carbonic Anhydrase (STPCA2-1) | Mummadisetti et al., 2021; Moya et al., 2008; Bertucci et al., 2011 |
| Gene g3745.t1 | Annotated: CARP4 | Mummadisetti et al., 2021; Puverel et al., 2005; Mass et al., 2013; Ramos-Silva et al., 2013; Mas et al., 2016; Takeuchi et al., 2016; Neder et al., 2019; Peled et al., 2020 |
| Gene: 10186 | Annotated: Protocadherin (PC2) | Mummadisetti et al., 2021; Drake et al., 2013; Takeuchi et al., 2016 |
| Gene: g30 | Annotated: Protoacadherin (PC4) | Mummadisetti et al., 2021; Drake et al., 2013; Takeuchi et al., 2016 |
| Gene: g10186 | Annotated: CARP4 | Mummadisetti et al., 2021; Puverel et al., 2005; Mass et al., 2013; Ramos-Silva et al., 2013; Mas et al., 2016; Takeuchi et al., 2016; Neder et al., 2019; Peled et al., 2020 |
| Gene: g10187 | Annotated: Protocadherin (PC3) | Mummadisetti et al., 2021; Drake et al., 2013; Takeuchi et al., 2016 |
| Gene: g10188 | Annotated: Protocadherin (PC3) | Mummadisetti et al., 2021; Drake et al., 2013; Takeuchi et al., 2016 |
| Gene: g11190 | Annotated: USOMP12 | Mummadisetti et al., 2021 |
| Gene: g13552 | Acidic SOMP (Full-Length p27) | Mummadisetti et al., 2021 |
| Gene: g14733 | Annotated: CARP4 | Mummadisetti et al., 2021; Puverel et al., 2005; Mass et al., 2013; Ramos-Silva et al., 2013; Mas et al., 2016; Takeuchi et al., 2016; Neder et al., 2019; Peled et al., 2020 |
| Gene: g1484 | Annotated: CARP1 | Mummadisetti et al., 2021; Mass et al., 2013; Neder et al., 2019 |
| Gene: g15294.t1 | Annotated: Vitellogenin | Mummadisetti et al., 2021; Drake et al., 2013; Peled et al., 2020 |
| Gene: g15955 | Annotated: MAM and LDL receptor-containing protein (MAM LDL-2) | Mummadisetti et al., 2021; Drake et al., 2013; Ramos-Silva et al., 2013; Takeuchi et al., 2016; Peled et al., 2020 |
| Gene: g1647 | Annotated: MAM and LDL receptor-containing (MAM LDL-1) | Mummadisetti et al., 2021; Drake et al., 2013; Ramos-Silva et al., 2013; Takeuchi et al., 2016; Peled et al., 2020 |
| Gene: g2115 | Annotated: Cadherin | Mummadisetti et al., 2021; Drake et al., 2013; Takeuchi et al., 2016 |
| Gene: g2116 | Annotated: Protocadherin (PC1) | Mummadisetti et al., 2021; Drake et al., 2013; Takeuchi et al., 2016 |
| Gene: g22569 | Annotated: Fibronectin | Mummadisetti et al., 2021 |
| Gene: g24177 | Annotated: Protocadherin (PC5) | Mummadisetti et al., 2021; Drake et al., 2013; Takeuchi et al., 2016 |
| Gene: g27814 | Annotated: carbonic anhydrase (STPCA2-2) | Mummadisetti et al., 2021 |
| Gene: g2829 | Annotated: Thrombospondin-like protein (Thrombospondin) | Mummadisetti et al., 2021 |
| Gene: g2829.t1 | Annotated: Coadhesin | Mummadisetti et al., 2021; Drake et al., 2013; Peled et al., 2020 |
| Gene: g29034.t1 | Annotated: CarbonicAnhyrase | Mummadisetti et al., 2021; Moya et al., 2008; Bertucci et al., 2011 |
| Gene: g30385.t1 | Annotated: USOMPS13 | Mummadisetti et al., 2021 |
| Gene: g34749 | Annotated: EGF and LamininG-Like (EGF LamG2) | Mummadisetti et al., 2021; Ramos-Silva et al., 2013; Takeuchi et al., 2016; Peled et al., 2020 |
| Gene: g37058 | Annotated: Fibronectin (Fibronectin-2) | Mummadisetti et al., 2021 |
| Gene: g38128 | Annotated: α-Collagen | Mummadisetti et al., 2021; Drake et al., 2013; Peled et al., 2020 |
| Gene: g38881 | Annotated: Uncharacterized protein (USOMP14) | Mummadisetti et al., 2021 |
| Gene: g39770 | Annotated: Kielin-Like | Mummadisetti et al., 2021 |
| Gene: g5735.t1 | Annotated: Tolloid-Like | Mummadisetti et al., 2021 |
| Gene: g7086 | Annotated: EGF and LamininG-Like (EGF LamG1) | Mummadisetti et al., 2021; Drake et al., 2013; Ramos-Silva et al., 2013; Takeuchi et al., 2016; Peled et al., 2020 |
| Gene: g7193 | Annotated: CARP2 | Mummadisetti et al., 2021 |
| Gene: g8396 | Annotated: N/A, named it CARP6-partial | Mummadisetti et al., 2021 |
| Gene: g907 | Annotated: Zona Pellucida (ZP domain-containing) | Mummadisetti et al., 2021; Drake et al., 2013; Takeuchi et al., 2016; Peled et al., 2020 |
| Gene: g9094 | Annotated: Actin | Mummadisetti et al., 2021 |
| Gene: Locus_65031_Transcript_1/1 | Annotated: CARP5 | Mummadisetti et al., 2021 |
| Gene: Locus_8768_Transcript_1/2 | Annotated: Sushi-domain | Mummadisetti et al., 2021 |
| AGE35225.2 | CARP1 | Mass et al., 2013; Mass et al., 2016; Neder et al., 2019 |
| AGE35226.1 | CARP3 | Mass et al., 2013; Mass et al., 2016; Neder et al., 2019 |
| AGE35227.1 | CARP2 | Mass et al., 2013; Mass et al., 2016; Neder et al., 2019 |
| ACE95141.1 | carbonic anhydrase | Moya et al., 2008 |
| PFX12726.1 | Retrovirus-related Pol polyprotein from transposon 17.6 | Peled et al., 2020 |
| PFX12813.1 | hypothetical protein AWC38_SpisGene23165 | Peled et al., 2020 |
| PFX13778.1 | Sacsin | Peled et al., 2020 |
| PFX14205.1 | Proto-oncogene tyrosine-protein kinase receptor Ret, partial | Peled et al., 2020 |
| PFX15740.1 | Protein FAM208A | Peled et al., 2020 |
| PFX16398.1 | hypothetical protein AWC38_SpisGene19330 | Peled et al., 2020 |
| PFX18785.1 | Mucin-4 | Peled et al., 2020; Takeuchi et al., 2016 |
| PFX26597.1 | Complement C3 | Peled et al., 2020 |
| PFX26751.1 | Transmembrane protease serine 9 | Peled et al., 2020 |
| PFX27832.1 | Poly [ADP-ribose] polymerase 11 | Peled et al., 2020 |
| PFX30831.1 | hypothetical protein AWC38_SpisGene4366 | Peled et al., 2020 |
| PFX30903.1 | hypothetical protein AWC38_SpisGene4292 | Peled et al., 2020 |
| PFX31810.1 | Nidogen-2 | Peled et al., 2020 |
| XP_022778254.1 | uncharacterized protein LOC111319781 | Peled et al., 2020 |
| XP_022778283.1 | uncharacterized protein LOC111319816, partial | Peled et al., 2020 |
| XP_022779720.1 | vitellogenin-like | Peled et al., 2020; Drake et al., 2013 |
| XP_022780303.1 | LOW QUALITY PROTEIN: uncharacterized protein LOC111321626 | Peled et al., 2020 |
| XP_022780690.1 | skeletal aspartic acid-rich protein 2-like (CARP5) | Peled et al., 2020; Ramos-Silva et al., 2013; Takeuchi et al., 2016 |
| XP_022780694.1 | CUB and peptidase domain-containing protein 2-like | Peled et al., 2020 |
| XP_022782398.1 | skeletal aspartic acid-rich protein 1-like (CARP4) | Peled et al., 2020; Puverel et al., 2005; Mass et al., 2013; Ramos-Silva et al., 2013; Mass et al., 2016; Takeuchi et al., 2016; Neder et al., 2019 |
| XP_022783044.1 | uncharacterized protein LOC111323869 | Peled et al., 2020 |
| XP_022783323.1 | phosphopantothenoylcysteine decarboxylase subunit VHS3-like | Peled et al., 2020 |
| XP_022783415.1 | coadhesin-like isoform X3 | Peled et al., 2020; Drake et al., 2013 |
| XP_022783952.1 | collagenase 3-like | Peled et al., 2020; Drake et al., 2013 |
| XP_022784623.1 | cation channel sperm-associated protein subunit beta-like | Peled et al., 2020 |
| XP_022786582.1 | synapsin-2-like isoform X2 | Peled et al., 2020 |
| XP_022786918.1 | major yolk protein-like isoform X2 | Peled et al., 2020; Drake et al., 2013 |
| XP_022788227.1 | hephaestin-like protein | Peled et al., 2020; Takeuchi et al., 2016 |
| XP_022788730.1 | chymotrypsin-like elastase family member 1 | Peled et al., 2020 |
| XP_022789591.1 | endothelin-converting enzyme 1-like isoform X2 | Peled et al., 2020 |
| XP_022789932.1 | MAGUK p55 subfamily member 7-like | Peled et al., 2020 |
| XP_022790441.1 | PHD finger protein 21A-like | Peled et al., 2020 |
| XP_022792212.1 | ras-like protein 3 | Peled et al., 2020 |
| XP_022794122.1 | galaxin-like isoform X2 | Peled et al., 2020; Ramos-Silva et al., 2013; Takeuchi et al., 2016 |

| Accession number/Gene ID | Definition | References |
|---|---|---|
| XP_022794351.1 | mammalian ependymin-related protein 1-like | Peled et al., 2020 |
| XP_022794736.1 | MAM and LDL-receptor class A domain-containing protein 2-like | Peled et al., 2020 |
| XP_022796981.1 | uncharacterized skeletal organic matrix protein 8-like | Peled et al., 2020 |
| XP_022796982.1 | uncharacterized protein LOC111335364 | Peled et al., 2020 |
| XP_022798902.1 | low-density lipoprotein receptor-related protein 8-like | Peled et al., 2020 |
| XP_022799089.1 | CUB domain-containing protein-like isoform X2 | Peled et al., 2020 |
| XP_022799541.1 | uncharacterized protein LOC111337489 | Peled et al., 2020 |
| XP_022803524.1 | digestive cysteine proteinase 1-like | Peled et al., 2020 |
| XP_022803808.1 | deleted in malignant brain tumors 1 protein-like | Peled et al., 2020 |
| XP_022803872.1 | spore wall protein 2-like isoform X3 | Peled et al., 2020 |
| XP_022803894.1 | uncharacterized protein LOC111341206 | Peled et al., 2020 |
| XP_022804012.1 | EGF and laminin G domain-containing protein-like | Peled et al., 2020; Ramos-Silva et al., 2013; Takeuchi et al., 2016 |
| XP_022804785.1 | thioredoxin reductase 1, cytoplasmic-like | Peled et al., 2020 |
| XP_022805470.1 | uncharacterized protein LOC111342641 | Peled et al., 2020 |
| XP_022806326.1 | ZP domain-containing protein-like | Peled et al., 2020; Drake et al., 2013; Takeuchi et al., 2016 |
| XP_022806664.1 | protein lingerer-like | Peled et al., 2020 |
| XP_022806928.1 | SLIT-ROBO Rho GTPase-activating protein 1-like | Peled et al., 2020 |
| XP_022807143.1 | condensin-2 complex subunit D3-like | Peled et al., 2020 |
| XP_022807256.1 | uncharacterized protein LOC111344300 | Peled et al., 2020 |
| XP_022807807.1 | uncharacterized protein LOC111344812 | Peled et al., 2020 |
| XP_022808163.1 | uncharacterized protein LOC111345150 | Peled et al., 2020 |
| XP_022808576.1 | uncharacterized protein LOC111345553 isoform X2 | Peled et al., 2020 |
| XP_022809269.1 | microtubule-associated tumor suppressor 1 homolog isoform X1 | Peled et al., 2020 |
| XP_022809270.1 | microtubule-associated tumor suppressor 1 homolog isoform X2 | Peled et al., 2020 |
| XP_022810585.1 | von Willebrand factor D and EGF domain-containing protein-like | Peled et al., 2020 |
| JR970990.1 | CUB and peptidase domain-containing protein 1 | Ramos-Silva et al., 2013 |
| JR971508.1 | Uncharacterized skeletal organic matrix protein-6 (USOMP6) | Ramos-Silva et al., 2013 |
| JR972076.1 | Acidic skeletal organic matrix protein (Acidic SOMP) | Ramos-Silva et al., 2013 |
| JR973117.1 | Uncharacterized skeletal organic matrix protein-5 (USOMP-5) | Ramos-Silva et al., 2013 |
| JR976690.1 | Galaxin 2 | Ramos-Silva et al., 2013 |
| JR978035.1 | Ectin | Ramos-Silva et al., 2013 |
| JR980881.1 | EGF and laminin G domain-containing protein | Ramos-Silva et al., 2013 |
| JR982706.1 | Uncharacterized skeletal organic matrix protein-2 (USOMP-2) | Ramos-Silva et al., 2013 |
| JR983041.1 | Secreted acidic protein 2 (Amil-SAP2) | Ramos-Silva et al., 2013 |
| JR983175.1 | Glu-rich protein | Ramos-Silva et al., 2013 |
| JR986059.1 | Cephalotoxin-like protein | Ramos-Silva et al., 2013 |
| JR987773 | Mucin-like | Ramos-Silva et al., 2013 |
| JR989025.1 | CUB domain-containing protein | Ramos-Silva et al., 2013 |
| JR991083.1 | Collagen alpha-1 chain | Ramos-Silva et al., 2013 |
| JR991141.1 | polycystic kidney disease 1-related skeletal organic matrix protein (PKD1-related protein) | Ramos-Silva et al., 2013 |
| JR991407.1 | Skeletal acidic Asp-rich Protein 2 (SAARP2) | Ramos-Silva et al., 2013 |
| JR993827.1 | Neuroglian-like | Ramos-Silva et al., 2013 |
| JR994474.1 | MAM and LDL-receptor domain- containing protein 2 | Ramos-Silva et al., 2013; Drake et al., 2013 |
| JR997000.1 | Uncharacterized skeletal organic matrix protein-3 (USOMP-3) | Ramos-Silva et al., 2013 |
| JR998014.1 | Putative carbonic anhydrase | Ramos-Silva et al., 2013; Bertucci et al., 2011 |
| JR998260.1 | Uncharacterized skeletal organic matrix protein-7 (USOMP7) | Ramos-Silva et al., 2013 |
| JT001945.1 | Skeletal acidic Asp-rich Protein 1 (SAARP 1) | Ramos-Silva et al., 2013 |
| JT004498.1 | Uncharacterized skeletal organic matrix protein-4 (USOMP-4) | Ramos-Silva et al., 2013 |
| JT008002.1 | CUB and peptidase domain-containing protein 2 | Ramos-Silva et al., 2013 |
| JT011093.1 | Protocadherin-like | Ramos-Silva et al., 2013 |
| JT011118.1 | MAM and LDL-receptor domain- containing protein 1 | Ramos-Silva et al., 2013; Drake et al., 2013 |
| JT013217.1 | MAM and fibronectin- containing protein | Ramos-Silva et al., 2013 |
| JT013896.1 | Threonine-rich protein | Ramos-Silva et al., 2013 |
| JT014391.1 | Uncharacterized skeletal organic matrix protein-8 (USOMP-8) | Ramos-Silva et al., 2013 |
| JT016410.1 | MAM and fibronectin containing protein 2 | Ramos-Silva et al., 2013 |
| JT016638.1 | Coadhesin | Ramos-Silva et al., 2013 |
| JT018094.1 ; JT006291.1 | Secreted acidic protein 1 (Amil-SAP1) | Ramos-Silva et al., 2013 |
| JT019463.1 | Hephaestin-like | Ramos-Silva et al., 2013 |
| JT021412.1 | Uncharacterized skeletal organic matrix protein-1 (USOMP-1) | Ramos-Silva et al., 2013 |
| HM163215 | Acropora millepora galaxin | Reyes-Bermudez et al., 2009 |
| aug_v2a.14283.t1 | CUB domain-containing protein 2 | Takeuchi et al., 2016 |
| adi_EST_assem_30005 | CUB domain-containing protein 2 N-terminus | Takeuchi et al., 2016 |
| aug_v2a.09226.t1 | Adi-SAP2 | Takeuchi et al., 2016 |
| aug_v2a.00002.t1 | EP-like 2 | Takeuchi et al., 2016 |
| aug_v2a.01440.t1 (aug_v2a.01441.t1) | Adi-SAARP2 | Takeuchi et al., 2016; Ramos-Silva et al., 2013 |
| aug_v2a.02662.t1 | USOMP-1b | Takeuchi et al., 2016 |
| aug_v2a.02663.t1 | USOMP-1c | Takeuchi et al., 2016 |
| aug_v2a.05945.t1 | TSP-1 and VWA domain-containing | Takeuchi et al., 2016 |
| aug_v2a.06327.t1 | SAARP3 | Takeuchi et al., 2016; Ramos-Silva et al., 2013 |
| aug_v2a.06593.t1 | SAP1 | Takeuchi et al., 2016 |
| aug_v2a.07084.t1 | USOMP-10 | Takeuchi et al., 2016 |
| aug_v2a.08856.t1 | Vitellogenin-like | Takeuchi et al., 2016 |
| aug_v2a.09809.t1 | Mucin4-like protein | Takeuchi et al., 2016 |
| aug_v2a.09968.t1 | MAM and LDLr domain-containing protein | Takeuchi et al., 2016; Drake et al., 2013; Ramos-Silva et al., 2013 |
| aug_v2a.09969.t1 | MAM and LDLr domain-containing protein | Takeuchi et al., 2016; Drake et al., 2013; Ramos-Silva et al., 2013 |
| aug_v2a.10941.t1 | CUB domain-containing protein 1 | Takeuchi et al., 2016 |
| aug_v2a.11068.t1 | SAARP1 | Takeuchi et al., 2016; Ramos-Silva et al., 2013 |
| aug_v2a.15064.t1 | Cystein-rich | Takeuchi et al., 2016 |
| aug_v2a.15065.t1 | galaxin2 | Takeuchi et al., 2016; Ramos-Silva et al., 2013 |
| aug_v2a.15580.t1 | Laminin G domain-containing protein | Takeuchi et al., 2016; Ramos-Silva et al., 2013 |
| aug_v2a.18376.t1 | USOMP-11 | Takeuchi et al., 2016 |
| aug_v2a.18631.t1 | galaxin | Takeuchi et al., 2016; Ramos-Silva et al., 2013 |
| aug_v2a.20893.t1 | USOMP-9 | Takeuchi et al., 2016 |
| aug_v2a.21723.t1 | USOMP-1a | Takeuchi et al., 2016 |
| aug_v2a.22918.t1 | EP-like1 | Takeuchi et al., 2016 |
| aug_v2a.02830.t1 | polycystic kidney disease 1-related (PKD1-related) protein | Takeuchi et al., 2016 |
| aug_v2a.06122.t1 | EGF and laminin G domain-containing protein | Takeuchi et al., 2016; Ramos-Silva et al., 2013 |
| aug_v2a.06123.t1 | EGF and laminin G domain-containing protein | Takeuchi et al., 2016; Ramos-Silva et al., 2013 |
| aug_v2a.07627.t1 | Zona pellucida domain-containing protein | Takeuchi et al., 2016; Drake et al., 2013 |
| aug_v2a.19518.t1 | Protocadherin-like | Takeuchi et al., 2016 |
| aug_v2a.24015.t1 | Hephaestin-like protein | Takeuchi et al., 2016 |
| aug_v2a.24512.t1 | EGF and laminin G domain-containing protein | Takeuchi et al., 2016; Ramos-Silva et al., 2013 |
| AAD11470.1 | L-type calcium channel alpha-1 subunit | Zoccola et al., 1999 |
| AAR13013.1 | plasma membrane calcium ATPase | Zoccola et al., 2004 |
| AJQ31790.1 | solute carrier family 4 member gamma | Zoccola et al., 2015 |
| XP_022791567.1 | anion exchange protein 2-like | Zoccola et al., 2015 |
| XP_022800320.1 | sulfate anion transporter 1-like isoform X1 | Zoccola et al., 2015 |
| XP_022800339.1 | sulfate anion transporter 1-like isoform X1 | Zoccola et al., 2015 |
| XP_022801463.1 | sodium bicarbonate cotransporter 3-like isoform X2 | Zoccola et al., 2015 |
| XP_022781731.1 | sodium-independent sulfate anion transporter-like 2C transcript variant X1 | Zoccola et al., 2015 |
| XP_022783031.1 | sodium bicarbonate transporter-like protein 11 | Zoccola et al., 2015 |
| XP_022788270.1 | band 3 anion transport protein-like | Zoccola et al., 2015 |
| XP_022800329.1 | sulfate anion transporter 1-like isoform X2 | Zoccola et al., 2015 |

Table S5. Photosynthetic parameters of algal endosymbiont. Parameters measured with a MAXI Imaging PAM, photosystem II (PSII) maximal quantum yield ($F_v/F_m$), maximum values of non-photochemical quenching ($NPQ_{max}$), maximum values of relative electron transport rate ($rETR_{max}$), initial slope (α) and minimal photoinhibition point ($E_k$). Values are represented as means ± SEM. Asterisk indicates significant differences between treatments (N = 9 for each, P value < 0.05).

| **Parameter** | **pH 8.2** | **pH 7.8** | **pH 7.6** |
|---|---|---|---|
| Fv/Fm* | 0.58 ± 0.02 | 0.59 ± 0.01 | 0.62 ± 0.004 |
| $NPQ_{max}$ | 0.68 ± 0.12 | 0.67 ± 0.03 | 0.73 ± 0.03 |
| $rETR_{max}$* | 56.65 ± 6.16 | 54.67 ± 2.39 | 75.02 ± 4.86 |
| α | 0.92 ± 0.05 | 0.97 ± 0.04 | 0.80 ± 0.05 |
| $E_k$ | 66.56 ± 12.93 | 58.02 ± 4.54 | 98.85 ± 11.18 |

### Supplementary References


1. Shefy D, Shashar N, Rinkevich B. The reproduction of the Red Sea coral *Stylophora pistillata* from Eilat: 4-decade perspective. Mar Biol.165(2):27.

2. Goodbody-Gringley G, Wong KH, Becker DM, Glennon K, de Putron SJ. Reproductive ecology and early life history traits of the brooding coral, *Porites astreoides*, from shallow to mesophotic zones. Coral Reefs. 37(2):483–94.

3. Goodbody-Gringley G. Diel planulation by the brooding coral *Favia fragum* (Esper, 1797). J Exp Mar Biol Ecol. 2010;389(1–2):70–4.

4. Pierrot D, Lewis E, Wallace DWR. MS Excel program developed for CO2 system calculations. ORNLCDIAC-105a Carbon Dioxide Inf Anal Cent Oak Ridge Natl Lab US Dep Energy Oak Ridge Tenn. 2006;10.

15. Gilis M, Meibom A, Domart-Coulon I, Grauby O, Stolarski J, Baronnet A. Biomineralization in newly settled recruits of the scleractinian coral *Pocillopora*

*damicornis*: Biomineralization in *P. damicornis* recruits. J Morphol. 2014;275(12):1349–65.
